# Climatic niche differentiation between native and non-native ranges is widespread in Ponto-Caspian amphipods

**DOI:** 10.1101/2023.05.23.541880

**Authors:** Eglė Šidagytė-Copilas, Denis Copilaș-Ciocianu

**Author notes:** corresponding author, Nature Research Centre, Akademijos 2, 08412, Vilnius, Lithuania.

## Abstract

1. Niche conservatism posits that a species’ non-native populations establish in areas that match the native environmental conditions. Although the Ponto-Caspian region is a major source of inland aquatic alien species, the extent to which their climatic niches diverged between the invasive and native ranges remains poorly understood.
2. Using an n-dimensional hypervolume approach, we quantified climatic niche overlap and inferred patterns of niche differentiation (shift, contraction, or expansion) among native and invaded ranges for 12 widespread Ponto-Caspian amphipod species (six genera in three families).
3. Our results show that all investigated species experience substantially different climatic conditions in the invaded range. The invasive niche either contracted (five species), shifted (four species), expanded and shifted (two species), or shifted and contracted (one species) relative to the native niche.
4. We conclude that although the focal taxa share a common geographic origin and evolutionary history, they exhibit disparate patterns of climatic niche change outside the native range. The niche conservatism hypothesis receives mixed support given that half of the studied species underwent niche shifts/expansion. Furthermore, congeners exhibited both identical and contrasting patterns of niche differentiation, suggesting a limited phylogenetic effect.
5. The uncovered diversity of niche dynamics among closely related species indicates that each has a unique potential for invasiveness and long-term persistence. This has important implications for predicting invasion risk and refining management strategies.

## 1. INTRODUCTION

Biological invasions are fundamentally reshaping global biodiversity with significant ecological, socio-economic, and health consequences (Ehrenfeld, 2010; Simberloff *et al*., 2013; Ogden *et al*., 2019; Diagne *et al*., 2021; Sayol *et al*., 2021). At the same time, introduced species can be regarded as invaluable models for studying evolutionary and ecological processes at contemporary time scales (Suarez & Tsutsui, 2008; Prentis *et al*., 2008; Wiens *et al*., 2019).

One of the main factors that determine the outcome of non-native species (alien or invasive) establishment is the degree to which the non-native environment diverges from the native one. Niche conservatism posits that species usually tend to conserve their niches (Wiens & Graham, 2005). Consequently, alien species should establish in regions that exhibit similar climatic conditions to the native range, thereby retaining their parental niche. However, the evidence for climatic niche conservatism in non-native species is contradictory, with studies finding widespread evidence for either niche conservatism (Petitpierre *et al*., 2012; Liu *et al*., 2020) or niche divergence (Atwater, Ervine & Barney, 2017; Torres *et al*., 2018). This is apparently due to methodological and interpretational issues (Bates & Bertelsmeier, 2021). Nevertheless, it appears that climatic niches tend to be less conserved in aquatic species (Lauzeral *et al*., 2011; Torres *et al*., 2018; Liu *et al*., 2020; Jourdan, Riesch & Cunze, 2021).

The Ponto-Caspian region (the Black, Azov, Caspian, and Aral seas as well as adjacent lagoons and river mouths) harbors a unique endemic aquatic fauna (Cristescu & Hebert, 2005; Naseka & Bogutskaya, 2009; Wesselingh *et al*., 2019; Copilaş-Ciocianu & Sidorov, 2022) characterized by a broad environmental tolerance (Reid & Orlova, 2002; Šidagytė & Arbačiauskas, 2016; Paiva *et al*., 2018; Borza *et al*., 2018; Pavel *et al*., 2021; Dobrzycka-Krahel, Stepien & Nuc, 2023). The adaptability of these species coupled with numerous anthropogenic factors (shipping, canals, and deliberate introductions) has transformed the Ponto-Caspian area into a major source of invasive species throughout the inland waters of the northern hemisphere (Bij de Vaate *et al*., 2002; Soto *et al*., 2023; Copilaş-Ciocianu, Sidorov & Šidagytė-Copilas, 2023b). The spread of many of these species is still ongoing (Copilaş-Ciocianu & Šidagytė-Copilas, 2022), generating significant ecological and economical impacts (Dermott *et al*., 1998; Vanderploeg *et al*., 2002; Strayer, 2009; Arbačiauskas *et al*., 2017; Gaye-Siessegger *et al*., 2022).

Despite the high number of species, broad phylogenetic diversity (polychaetes, mollusks, crustaceans, and fishes), and widespread distribution of invasive Ponto-Caspian taxa, little is known about the degree to which their niches in the invaded range diverged relative to the native range. The evidence so far suggests that in general most taxa seem to occupy lower and narrower salinity ranges throughout the invaded region (Pauli & Briski, 2018), but very little is known with respect to climate. The single study conducted to date found a significant niche shift in the zebra mussel (*Dreissena polymorpha*) in both its European and North American invaded ranges (Gallardo, zu Ermgassen & Aldridge, 2013).

Among the Ponto-Caspian fauna, amphipod crustaceans are taxonomically and ecologically the most diverse organismal group (Copilaş-Ciocianu & Sidorov, 2022), with up to 40% of species being reported outside the native range due to human intervention (Copilaş-Ciocianu *et al*., 2023b). Their non-native spread is often accompanied by extinction of native amphipods and restructuring of communities (Dermott *et al*., 1998; Van Riel *et al*., 2006; Arbačiauskas, 2008; Grabowski *et al*., 2009; Soto *et al*., 2022). One species, the killer shrimp (*Dikerogammarus villosus*), is considered one of the worst 100 invasive species world-wide (Rewicz *et al*., 2014).

The non-native ranges of Ponto-Caspian amphipods vary drastically in size from local to intercontinental, while significant differences also exist throughout the native range (Copilaş-Ciocianu *et al*., 2023b). Additionally, some species have narrow native and broad non-native ranges, while others exhibit opposite patterns (Copilaş-Ciocianu *et al*., 2023b). This range-size disparity within and outside the native region might reflect the environmental tolerance of these species given that range-size reflects niche breadth (Slatyer, Hirst & Sexton, 2013; Sheth & Angert, 2014; Ficetola, Lunghi & Manenti, 2020), including in gammarid amphipods (Gaston & Spicer, 2001). Therefore, we hypothesize that Ponto-Caspian amphipods should display a broad dynamic of niche differentiation patterns between the native and non-native ranges. We tested this hypothesis by using an n-dimensional hypervolume approach on a curated dataset of ∼8000 records belonging to the 12 most widespread species.

## 2. METHODS

### 2.1 Data acquisition

All the occurrence data used in this study was compiled in our previous research (Copilaş-Ciocianu *et al*., 2023b) and is freely available on Figshare (https://doi.org/10.6084/m9.figshare.20984461.v1). This curated dataset contains 8145 distribution points from 39 species obtained from own research, literature (322 sources), and online repositories such as the Global Biodiversity Information Facility (https://www.gbif.org/). However, distribution data from both native and invasive ranges were available for 13 species. From these, we further excluded *Dikerogammarus bispinosus* since the available data from the native region is insufficient and most likely this taxon comprises three distinct species, two of which are undescribed and their ranges are poorly known (Morhun *et al*., 2022). The final dataset contained 12 species and 7794 records (Tables 1, S1). The native range was conservatively defined to encompass the Azov, Black and Caspian seas, their adjacent lagoons and estuaries/deltas (Wesselingh *et al*., 2019; Copilaş-Ciocianu & Sidorov, 2022; Copilaş-Ciocianu *et al*., 2023b). Areas outside this circumscription were considered non-native.

**Table 1.** Hypervolume-based climatic niche differentiation among native and invaded ranges of Ponto-Caspian amphipods. The β_total_ index was calculated based on Jaccard similarity. Values based on the Sørensen-Dice coefficient are presented in the supplementary Table S3.

| Species | Total native volume | Total invasive volume | Unique native volume | Unique invasive volume | Niche volume overlap | Jaccard similarity | Niche centroid distance | $\beta_{\text{total}}$ | $\beta_{\text{replacement}}$ (% total differentiation) | $\beta_{\text{richness}}$ (% total differentiation) | Inferred niche change |
| --- | --- | --- | --- | --- | --- | --- | --- | --- | --- | --- | --- |
| *Chaetogammarus ischnus* | 3528 | 2339 | 2909 | 1720 | 619 | 0.12 | 3.81 | 0.87 | 0.64 (74%) | 0.23 (26%) | shift |
| *Chaetogammarus warpachowskyi* | 437 | 314 | 357 | 234 | 80 | 0.12 | 5.67 | 0.88 | 0.69 (78%) | 0.18 (22%) | shift |
| *Chelicorophium curvispinum* | 2091 | 474 | 1780 | 162 | 311 | 0.14 | 4.24 | 0.86 | 0.14 (16%) | 0.72 (84%) | contraction |
| *Chelicorophium robustum* | 1414 | 204 | 1330 | 119 | 84 | 0.06 | 6.56 | 0.94 | 0.15 (16%) | 0.79 (84%) | contraction |
| *Chelicorophium sowinskyi* | 296 | 705 | 194 | 603 | 102 | 0.11 | 5.97 | 0.89 | 0.43 (48%) | 0.46 (52%) | expansion + shift |
| *Dikerogammarus haemobaphes* | 913 | 459 | 710 | 256 | 202 | 0.17 | 4.31 | 0.82 | 0.42 (51%) | 0.39 (48%) | shift + contraction |
| *Dikerogammarus villosus* | 216 | 684 | 131 | 599 | 84 | 0.10 | 5.49 | 0.89 | 0.32 (35%) | 0.58 (65%) | expansion + shift |
| *Obesogammarus crassus* | 5483 | 143 | 4500 | 447 | 982 | 0.17 | 3.9 | 0.83 | 0.14 (17%) | 0.69 (83%) | contraction |
| *Obesogammarus obesus* | 5956 | 1275 | 5497 | 816 | 459 | 0.07 | 4.46 | 0.93 | 0.23 (25%) | 0.69 (75%) | contraction |
| *Pontogammarus robustoides* | 7964 | 1180 | 7157 | 372 | 807 | 0.10 | 3.34 | 0.90 | 0.08 (9%) | 0.82 (91%) | contraction |
| *Pontogammarus sarsi* | 425 | 555 | 259 | 389 | 166 | 0.20 | 4.94 | 0.79 | 0.63 (80%) | 0.16 (20%) | shift |
| *Spirogammarus major* | 249 | 408 | 231 | 390 | 18 | 0.03 | 5.71 | 0.97 | 0.72 (74%) | 0.25 (26%) | shift |

Current climate data (∼1950–2000; 19 bioclimatic variables) were downloaded from the WorldClim database (https://www.worldclim.org/) (Hijmans *et al*., 2005) with DIVA GIS 7.5 (http://www.diva-gis.org/) at a 2.5-minute resolution. The full dataset is available in the supplementary Table S1. To reduce spatial sampling bias, we thinned the occurrence dataset of each species to eliminate points which were closer to one another than 5 km using the R package *spThin* v. 0.2.0 and 100 repetitions (Aiello-Lammens *et al*., 2015). After thinning, each species had a median of 175 (range = 45–1071) non-native records and a median of 35.5 (range = 9–79) native records. Details for each species are in the supplementary Table S2.

### 2.2 Data analysis

There are many methods for comparing climatic niches, all falling within four broad categories: hypervolumes, ordination, univariate, and ecological niche modelling (Fitzpatrick *et al*., 2007; Blonder *et al*., 2014; Guisan *et al*., 2014; Liu *et al*., 2020; Broennimann *et al*., 2021). Here we chose the hypervolume approach since, unlike the other methods, it defines niche space in more than two dimensions, thus being more realistic and in line with the Hutchinsonian multidimensional niche concept (Blonder, 2018), but it can also be sensitive to low sample size and high dimensionality (Qiao *et al*., 2017). Hypervolumes have been used extensively to compare climatic niches among species and populations (Mammola & Isaia, 2017; Pili *et al*., 2020; Zhang *et al*., 2020; Liu *et al*., 2020).

Using the R packages *FactoMineR* v. 2.8 (Lê, Josse & Husson, 2008; Husson *et al*., 2023) and *factoextra* v. 1.0.7 (Kassambra & Mundt, 2022), we ran and visualized a correlation-matrix-based principal component analysis (PCA) of all 19 bioclimatic variables for each species and retained the number of axes to encompass at least 80% of the variance. In such a way, the number of retained PCs varied between 3 and 4, depending on the species. We calculated the PCAs independently for each species since the bioclimatic parameters driving the niche changes could be species-specific and our aim was to specifically compare the niches of each species between the native and invaded ranges (i.e., absolute niche sizes are not comparable among species).

The obtained PCA axes were then used to construct (using kernel density estimation) and depict (sampling 10 000 random points) two hypervolumes for each species – each corresponding to the native and invasive climatic niches. This was done using the R package *hypervolume* v. 3.1.0 (Blonder *et al*., 2014, 2018, 2022). For each native-invasive hypervolume pair, we also extracted the n-dimensional volumes of each hypervolume, their overlap, and their unique parts, as well as the distance between centroids. We then input the hypervolume pairs into the R package *BAT* v. 2.9.2 (Cardoso, Rigal & Carvalho, 2015; Mammola & Cardoso, 2020) to estimate the β_total_ diversity index (based on the Jaccard similarity and the Sørensen–Dice coefficients) to quantify the total niche change on a scale from 0 (identical) to 1 (completely dissimilar) and its partitioning into the β_replacement_ (indicating niche shift) and β_richness_ (indicating either niche contraction or expansion) indices (Carvalho & Cardoso, 2020).

To aid with the grouping of the species according to the inferred climatic niche changes, we used hierarchical clustering. All the three β coefficients, the relative sizes of unique native range, unique invaded range, and their overlap were used as input into the distance matrix. This was done using default R functions and options.

## 3. RESULTS

In general, we find that the native climate for most species is drier, warmer, and with a more expressed seasonality than the non-native climate. However, there is a lot of heterogeneity among species (Fig. 1). As for the n-dimensional hypervolumes, all 12 investigated species exhibited very little overlap between the invasive and native climatic niches (Jaccard index = 0.06–0.17; β_total_ = 0.82– 0.97) (Table 1, Figs. S1–12). Invasive niche hypervolumes were either larger, similar, or smaller than the hypervolume of the native niche (Table 1, Fig. 2). We infer that *C. ischnus, C. warpachowskyi, P. sarsi*, and *S. major* experienced a niche shift as the contribution of the β_replacement_ index to the total differentiation (β_total_) ranged between 74 to 80% (Table 1, Fig. 2, S2–3, S11–12). Two species, i.e., *C. sowinskyi* and *D. villosus*, have likely undergone a niche expansion accompanied by a shift given that the contribution of β_richness_ (52 and 65%, respectively) was rather similar to that of β_replacement_ and the hypervolume of the invaded range was two to three time greater than that of the native one (Table 1, Fig. 2, S5, S7). For *D. haemobaphes*, we infer a niche contraction accompanied by a shift because both β_richness_ and β_replacement_ contributed equally to the total differentiation and the native hypervolume was almost twice greater than the invaded hypervolume (Table 1, Fig. 2, S6). The remaining five species (*C. curvipinum, C. robustum, O. crassus, O. obesus*, and *P. robustoides*) likely underwent a niche contraction given that the contribution of β_richness_ ranged between 75 to 91% and the invaded hypervolume was four to eight times smaller than the native one (Table 1, Fig. 2, S3, S4, S8-10). Using the Sørensen–Dice coefficient we obtained very similar results (Table S3).

**Figure 1.**
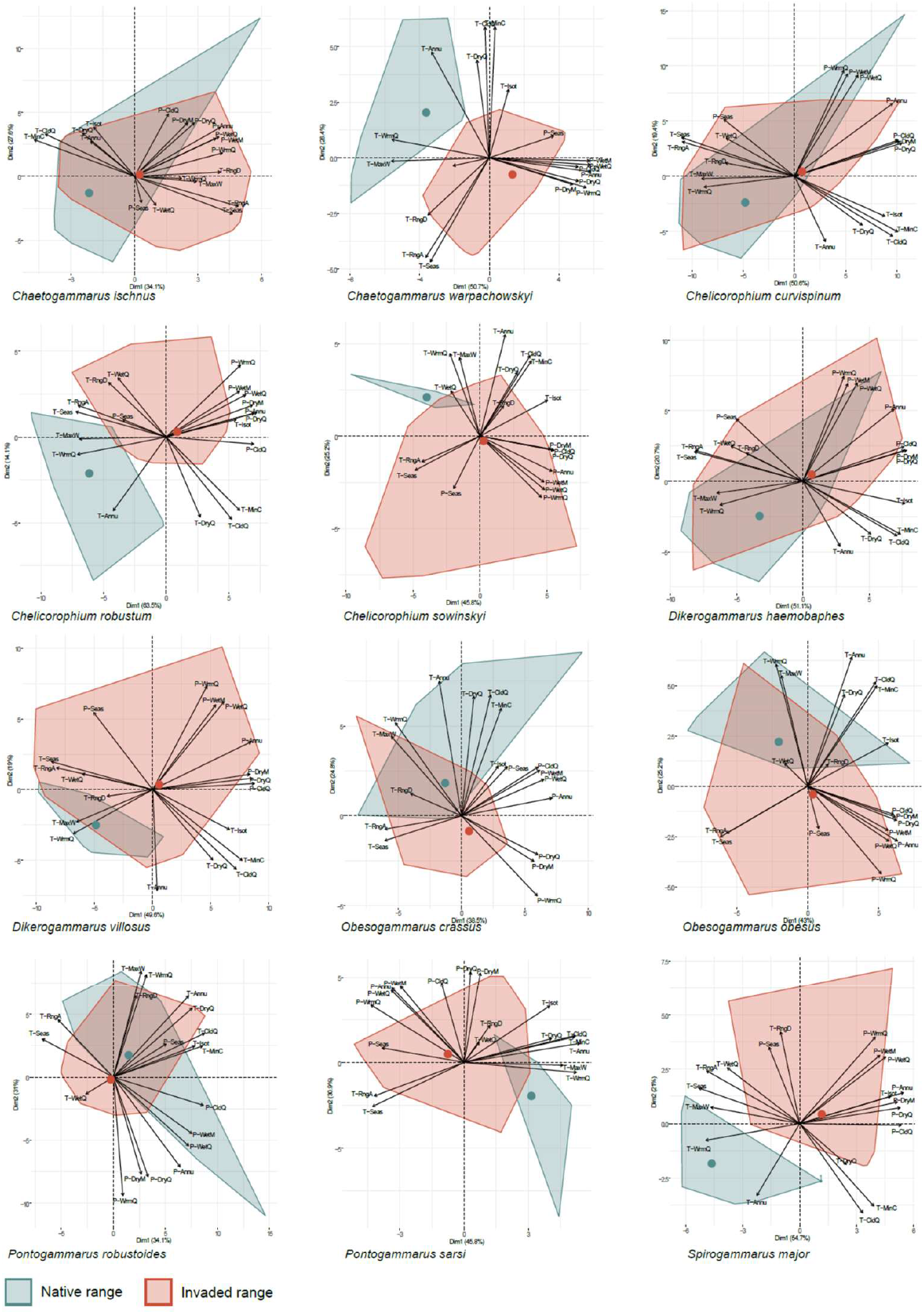
PCA scatterplots depicting differentiation among the climatic niches in the native (green) and invaded (red) ranges of 12 Ponto-Caspian amphipods. Colored dots indicate centroids. Abbreviations for the 19 bioclimatic variables are as follows: T-Annu – annual mean temperature; T-RngD – mean diurnal range; T-Isot – isothermality; T-Seas – temperature seasonality; T-MaxW – max temperature of warmest month; T-MinC – min temperature of coldest month; T-RngA – temperature annual range; T-WetQ – mean temperature of wettest quarter; T-DryQ – mean temperature of driest quarter; T-WrmQ – mean temperature of warmest quarter; T-CldQ – mean temperature of coldest quarter; P-Annu – annual precipitation; P-WetM – precipitation of wettest month; P-DryM – Precipitation of Driest Month; P-Seas – Precipitation Seasonality; P-WetQ – precipitation of wettest quarter; P-DryQ – precipitation of driest quarter; P-WrmQ – precipitation of warmest quarter; P-CldQ – precipitation of coldest quarter.

**Figure 2.**
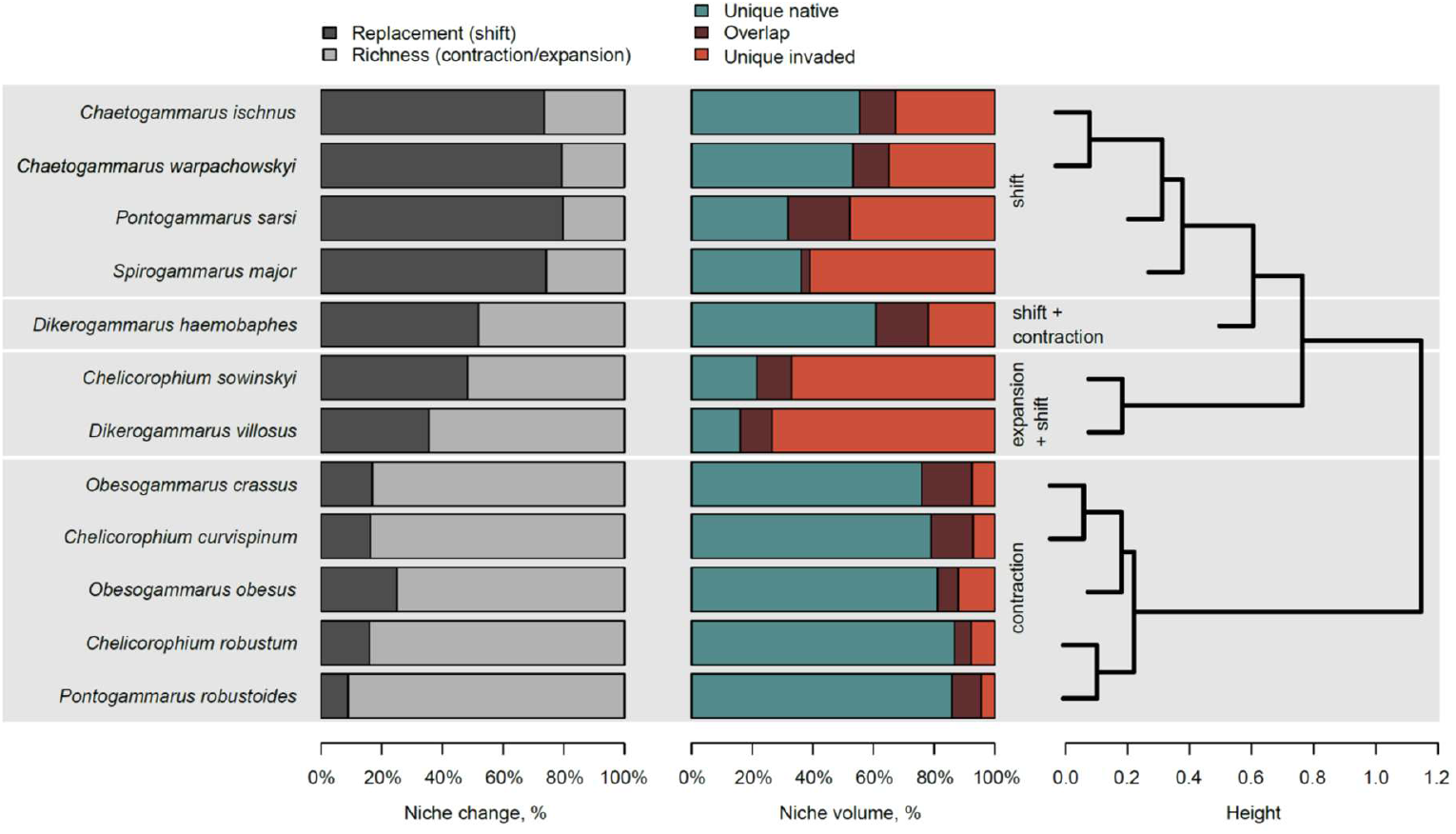
Climatic niche differentiation dynamics in alien Ponto-Caspian amphipods based on n-dimensional hypervolumes. Bar plots on the left show the relative contribution of the β_replacement_ (indicating niche shift) and the β_richness_ (indicating niche contraction/expansion) indices to the total differentiation (β_total_). Bar plots in the center show the relative sizes of the native and invasive climatic niches and their overlap. The dendrogram on the right indicates species clusters based on based on niche hypervolumes and niche differentiation indices.

## 4. DISCUSSION

### 4.1 Dynamics of niche differentiation

Our results reveal that climatic niche differentiation among native and invaded ranges is ubiquitous in all the 12 studied Ponto-Caspian amphipod species. On one hand, in half of the species we mostly detected a contraction of the invasive niche, which is in line with the niche conservatism hypothesis (Wiens & Graham, 2005; Liu *et al*., 2020). On the other hand, in the remaining half we find evidence for either niche shift (four species) or niche expansion accompanied by a shift (two species), supporting the view that niches might not be so conserved (Lauzeral *et al*., 2011; Atwater *et al*., 2017; Zhang *et al*., 2020; Jourdan *et al*., 2021). Therefore, overall, our study supports both views. The observed gradient in the β_replacement_ to β_richness_ ratio and native to invaded niche volume ratio without abrupt changes supports the existence of a continuum between no niche shift to a substantial niche shift. Thus, we agree with the view that focusing on the relative degree of niche change rather than a dichotomous classification paints a more realistic picture regarding niche differentiation (Bates & Bertelsmeier, 2021).

The uncovered pattern of niche differentiation dynamics is unexpectedly diverse, despite the common geographical origin and close evolutionary relationship between the focal species (Hou, Sket & Li, 2014; Copilaş-Ciocianu *et al*., 2022). Moreover, we find both homogeneous and heterogeneous patterns of niche differentiation within genera. For example, both studied *Chaetogammarus* species exhibit niche shifts while both *Obesogammarus* species exhibit niche contraction. Conversely, within *Chelicorophium, Dikerogammarus*, and *Pontogammarus* we find various, often contrasting (contraction–expansion) patterns of niche differentiation. Overall, this indicates a limited phylogenetic effect, highlighting the evolutionary lability of the climatic niche despite long term evolution in the geographically limited Ponto-Caspian region since the late Miocene (Copilaş-Ciocianu, Borko & Fišer, 2020).

We argue that the diversity of niche differentiation dynamics might be related to the broad climatic conditions present within the native range, which vary from humid subtropical through oceanic, and continental in the Black Sea region, and from Mediterranean through semi-arid and cold desert in the Caspian Sea region (Köppen climate classification; https://en.wikipedia.org/wiki/K%C3%B6ppen_climate_classification). Thus, many Ponto-Caspian species are exposed to a broad range of conditions, hence their propensity to invade outside the native range. This is further supported by the fact that all species that underwent a niche contraction are widespread in both the Black Sea and Caspian Sea basins, whereas the species that underwent a niche shift or expansion tend to be rarer or absent in the Caspian basin (Copilaş-Ciocianu *et al*., 2023b). Furthermore, the climate in the invaded range tends to be colder, wetter, and with a less pronounced seasonality than in the native range, thus possibly being more permeable to colonization given the apparent tolerance of these species to lower temperatures (Dobrzycka-Krahel, Kemp & Fidalgo, 2022). Similar findings were reported for the zebra mussel (Gallardo *et al*., 2013). It is likely that these patterns are common for many invasive Ponto-Caspian species given that their geographical ranges and invasion routes are similar to those of the amphipods (Jażdżewski, 1980; Bij de Vaate *et al*., 2002; Copilaş-Ciocianu *et al*., 2023b).

Future phylogeographic studies will help refine our understanding of niche differentiation dynamics by taking evolutionary lineages into account. The genetic evidence accrued so far indicates that some focal species are comprised of two mitochondrial lineages, each endemic to either the Black Sea or Caspian Sea drainages (Cristescu *et al*., 2004; Cristescu & Hebert, 2005; Rewicz *et al*., 2015; Jażdżewska *et al*., 2020; Copilaş-Ciocianu *et al*., 2022; Copilaş-Ciocianu *et al*., 2023a; Copilaş-Ciocianu *et al*., 2023). However, considering that their divergence is low and unconfirmed at the nuclear level, it is unclear whether these lineages are distinct evolutionary units equivalent to species. The only exception is *D. haemobaphes*, which harbors two highly divergent lineages in the Black Sea basin confirmed with nuclear and mitochondrial markers (Jażdżewska *et al*., 2020).

However, considering that both are sympatric and the new lineage has a narrow distribution, it is unlikely that this had an influence on the outcome of the analysis (Jażdżewska et al., 2020; Copilaş-Ciocianu, unpublished data).

### 4.2 Implications for management and conservation

Once invasive species establish throughout the non-native range, their eradication becomes impractical (Parkes & Panetta, 2009). Thus, predicting invasion is paramount. However, given that invasion risk prediction is based on niche conservatism, it may be problematic to anticipate the future spread of invasive species that undergo niche shifts or expansion (Petitpierre *et al*., 2012; Pili *et al*., 2020). It is unclear whether the Ponto-Caspian amphipods that predominantly experienced a niche expansion (*C. sowinskyi* and *D. villosus*) or a niche shift (*C. ischnus, C. warpachowskyi, P. sarsi*, and *S. major*) might be expected to further spread or not. This could be investigated using species distribution models that consider both the native and the invaded climate and future climate projections (Guisan *et al*., 2014; Eckert *et al*., 2020). Attempts made so far with *D. haemobaphes* and *D. villosus* have shown a limited future dispersal for both species (Cancellario *et al*., 2023), but this study did not consider potential niche shifts and inadequately represented the species in the native range (the extensive Caspian range of *D. haemobaphes* was omitted).

The six amphipod species that predominantly underwent a niche contraction might be expected to further spread until their invasive climatic niche breadth matches the native one. In these cases, species distribution models might prove invaluable in predicting invasion risk. This is especially important considering that on-going spread of Ponto-Caspian amphipods is routinely reported (Csabai *et al*., 2020; Lipinskaya *et al*., 2021; Copilaş-Ciocianu & Šidagytė-Copilas, 2022) and that the number of invasive crustaceans in Europe and temperate Asia are expected to rise substantially by 2050 (Seebens *et al*., 2021). However, some studies suggest that the likelihood of future invasions by Ponto-Caspian peracarid crustaceans remains low (Borza *et al*., 2017).

### 4.3 Methodological considerations

In this study we employed the approach of Carvalho and Cardoso (2020) for estimating niche shifts, contraction, and expansion. This method is somewhat analogous to the COUE scheme (Broennimann *et al*., 2012) where niche contraction is referred to as “unfilling” (ecological conditions unique to the native range and not present in the invaded range), niche overlap as “stability” (ecological conditions present in both native and invaded ranges), and niche expansion as “expansion” (ecological conditions unique to the invaded range and not present in the native range).

A key difference between the hypervolume approach employed in our study and the COUE scheme is that we used a species’ entire climatic niche space for calculating niche differentiation, whereas the latter uses climates that are analogous between the invaded and native range, while excluding non-analogous climates (Petitpierre *et al*., 2012; Guisan *et al*., 2014; Zhang *et al*., 2020). The reasoning behind using only analogous climates is either methodological (inability of species distribution models to extrapolate into non-analogous climates (Webber *et al*., 2011; Webber, Le Maitre & Kriticos, 2012; Pili *et al*., 2020)) or interpretational (inability to discern between fundamental or realized niche shifts when colonizing non-analogous climates (Petitpierre *et al*., 2012; Guisan *et al*., 2014)). However, it is also not possible to distinguish between fundamental or realized niche shifts in analogous climatic space as well because empirical distribution data cannot quantify a species fundamental niche (Soberón, 2007; Bates & Bertelsmeier, 2021). Only complementary experimental approaches can potentially determine the nature of these shifts (Pearman *et al*., 2008; Colautti & Lau, 2015; Bates & Bertelsmeier, 2021). The debate for including or excluding non-analogous climates is on-going, reflecting the need for methodological unification (Bates & Bertelsmeier, 2021).

## 5. CONCLUSIONS

We conclude that the most widespread Ponto-Caspian invasive amphipods experience substantially different climatic conditions throughout the non-native range relative to the native range. The uncovered diversity of niche differentiation dynamics is surprising given the close phylogenetic relationships and long-term evolution within the confines of the Ponto-Caspian realm. This indicates that each species has a distinct potential for invasiveness and long-term persistence. Ponto-Caspian amphipods may prove to be a useful model system for experimental studies focusing on the adaptive mechanisms of invasive species.

## Data availability

All the data used for analyses is available in the supplementary information.

## Supplementary tables

Table S1. Records and bioclimatic variables used for analyses.

Table S2. Number of records and basic statistics for each species before and after thinning with *spThin*.

Table S3. Measures of niche overlap and differentiation based on the Sørensen–Dice coefficient.

## Supplementary figures

Figs. S1–S12. Native and invasive niche hypervolumes for each of the 12 studies species. The large dots represent niche centroids. The small dots are 10,000 random points sampled from each hypervolume to delineate its shape and boundary.

