## Supplementary material for "Climatic niche differentiation between native and non-native ranges is widespread in Ponto-Caspian amphipods": Fig. S

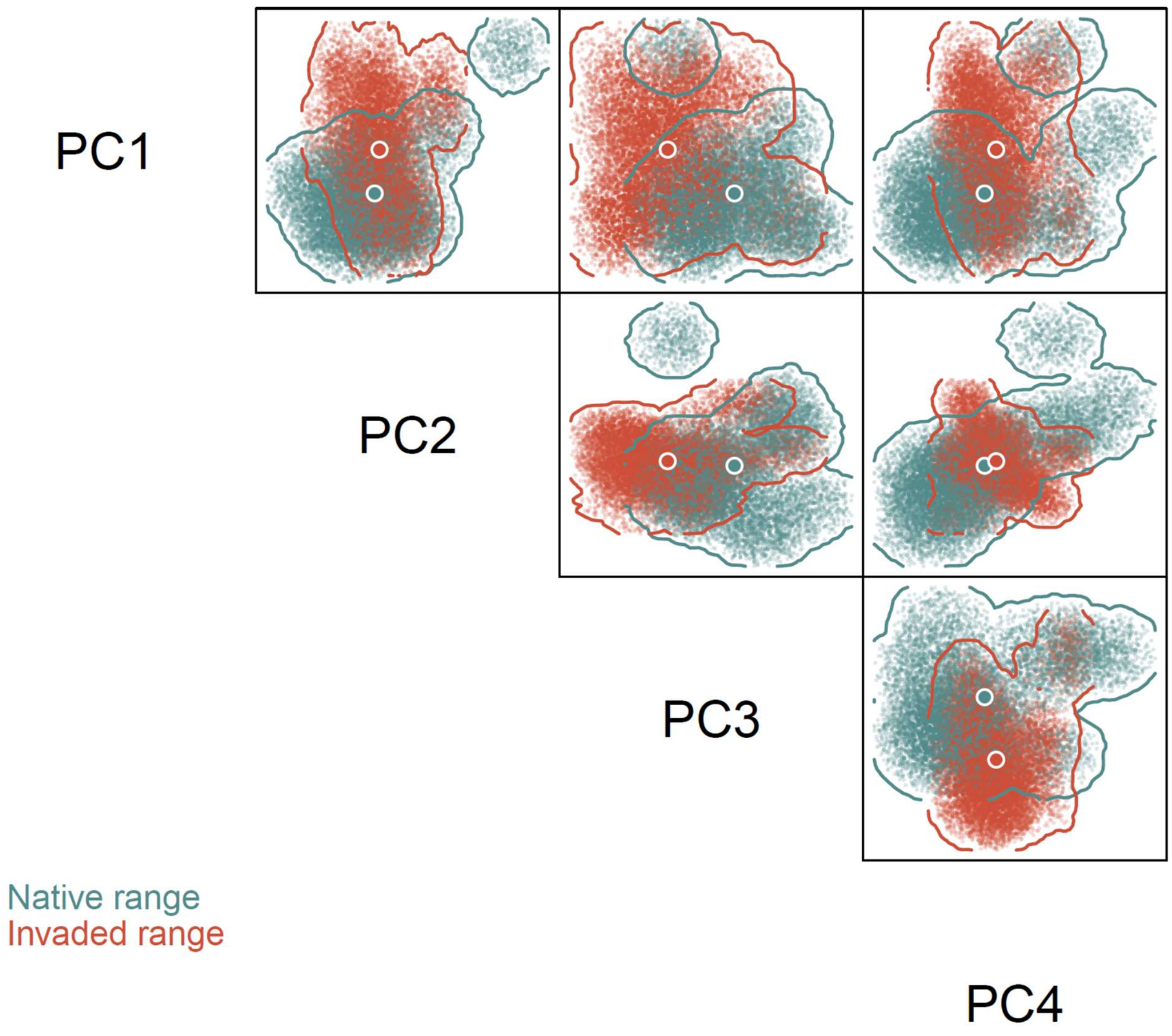


Fig. S1. Native and invasive niche hypervolumes of *Chaetogammarus ischnus*. The large dots represent niche centroids. The small dots are 10.000 random points sampled from each hypervolume to delineate its shape and boundary.

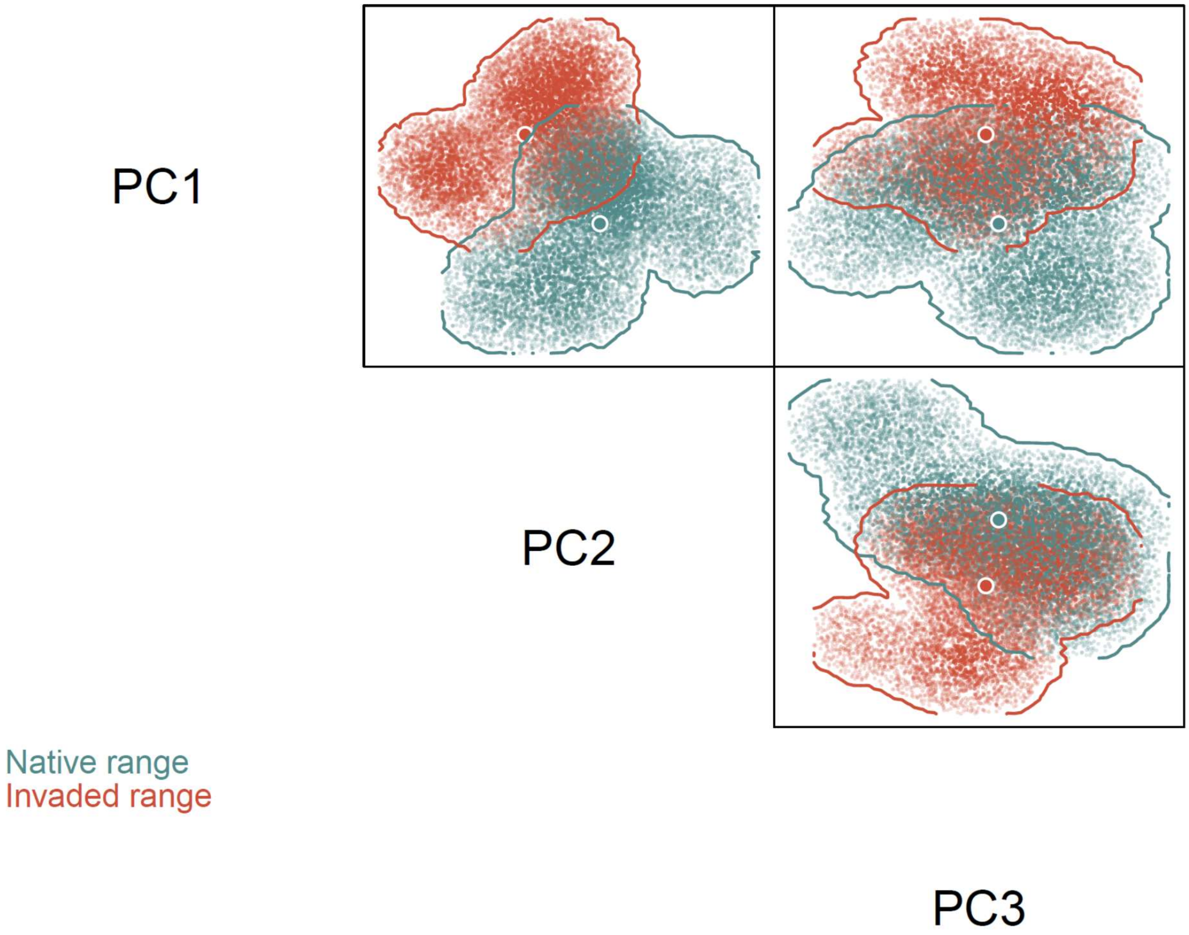


Fig. S2. Native and invasive niche hypervolumes of *Chaetogammarus warpachowskyi*. The large dots represent niche centroids. The small dots are 10.000 random points sampled from each hypervolume to delineate its shape and boundary.

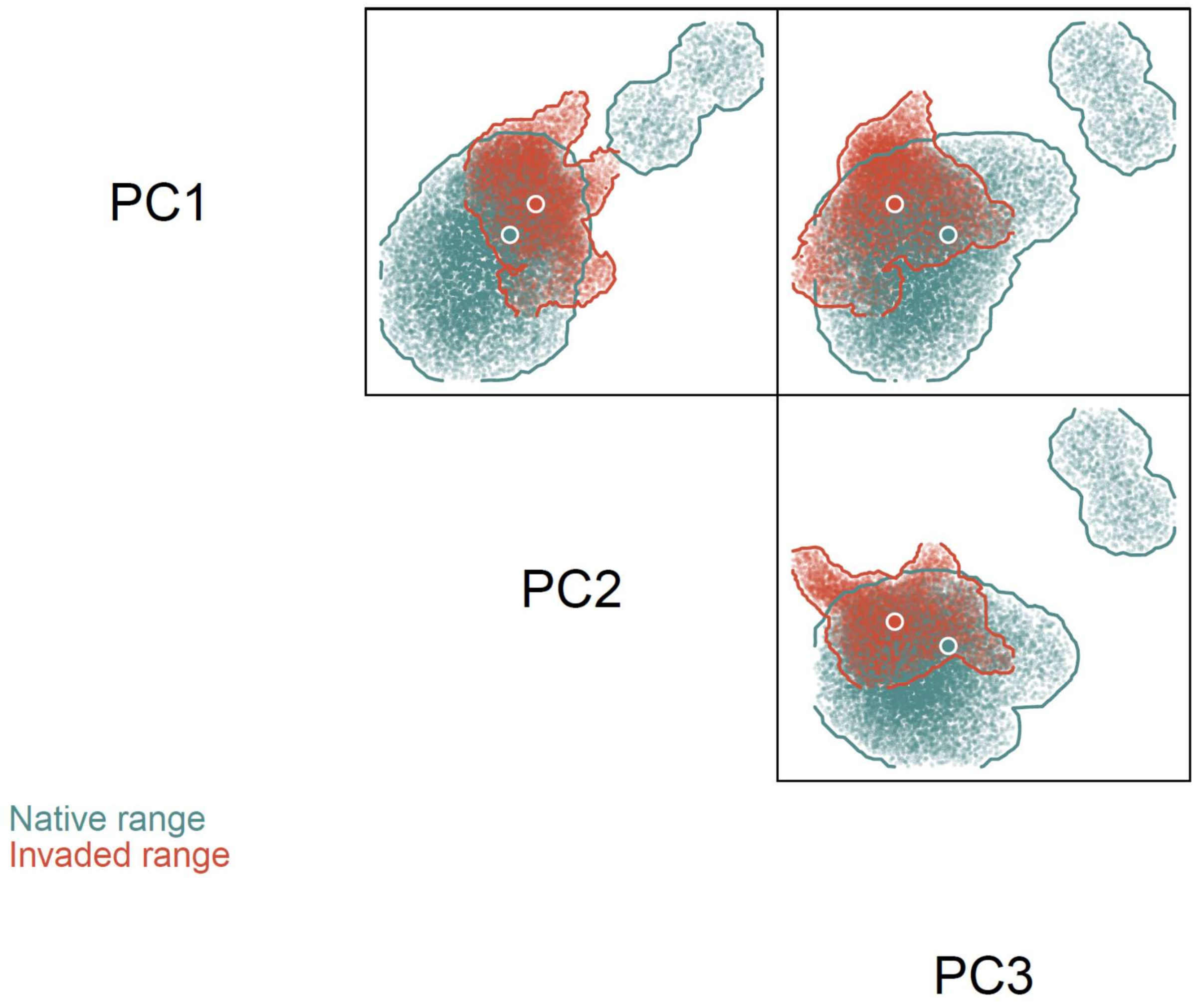


Fig. S3. Native and invasive niche hypervolumes of *Chelicorophium curvispinum*. The large dots represent niche centroids. The small dots are 10.000 random points sampled from each hypervolume to delineate its shape and boundary.

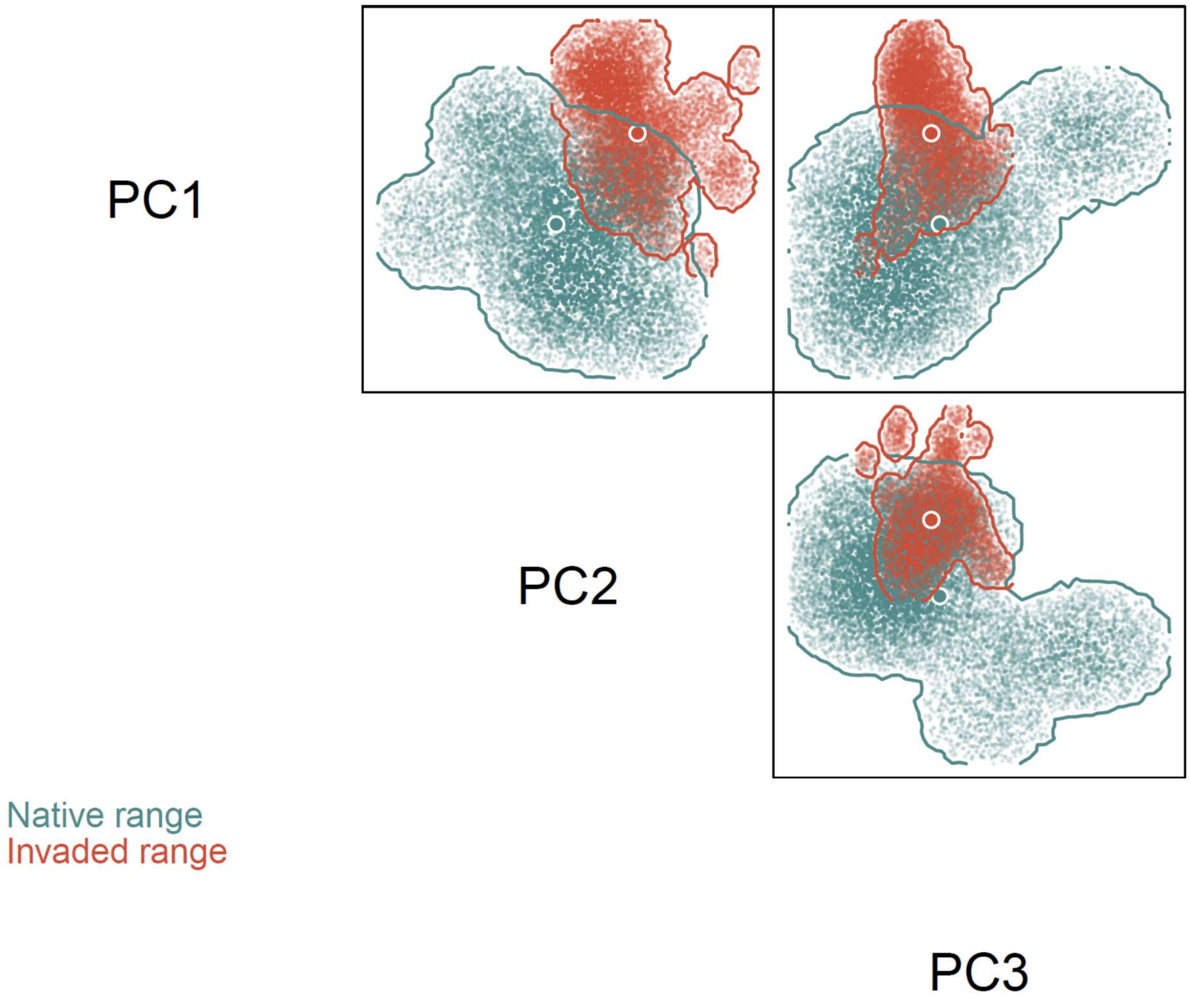


Fig. S4. Native and invasive niche hypervolumes of *Chelicorophium robustum*. The large dots represent niche centroids. The small dots are 10.000 random points sampled from each hypervolume to delineate its shape and boundary.

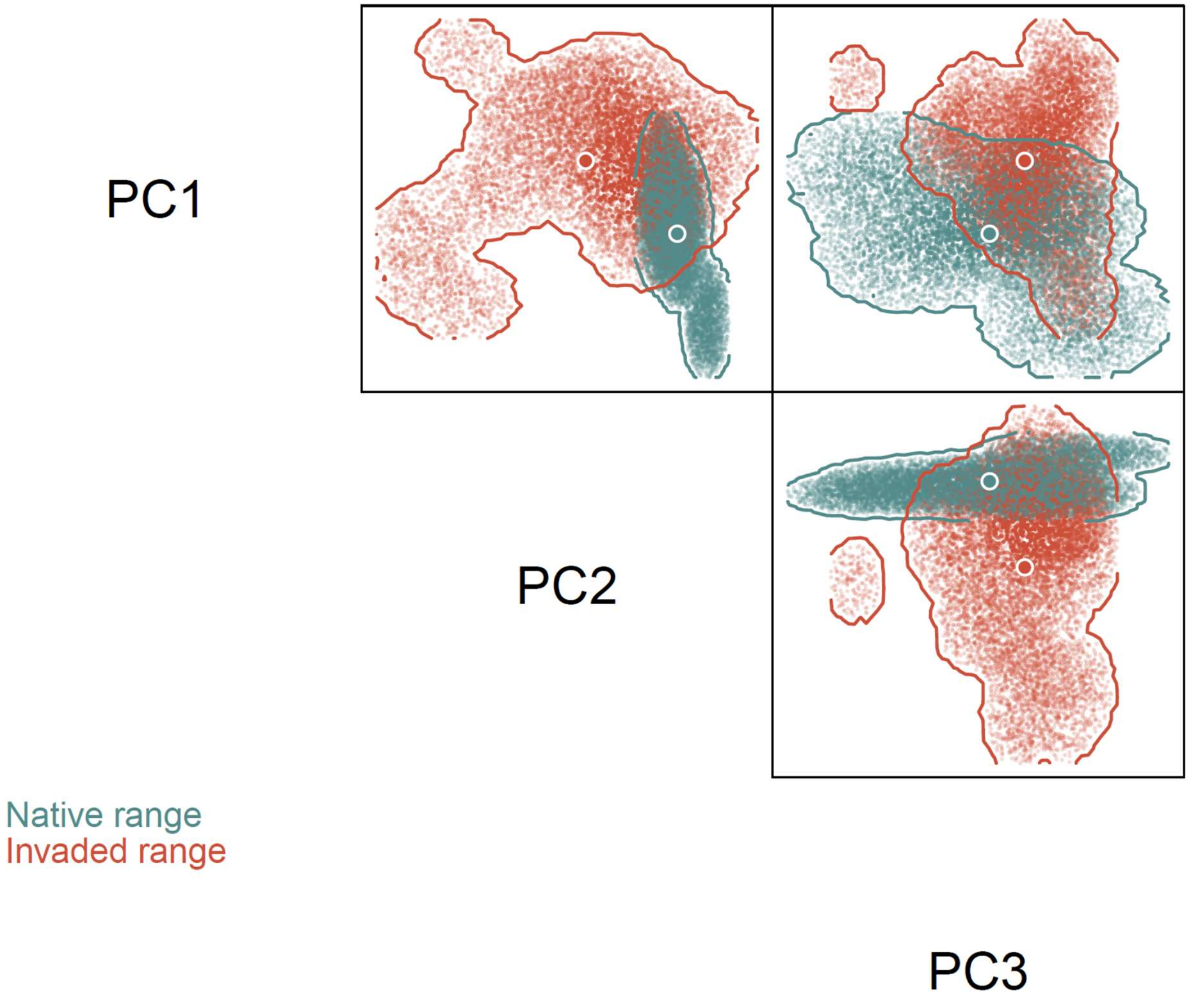


Fig. S5. Native and invasive niche hypervolumes of *Chelicorophium sowinskyi*. The large dots represent niche centroids. The small dots are 10.000 random points sampled from each hypervolume to delineate its shape and boundary.

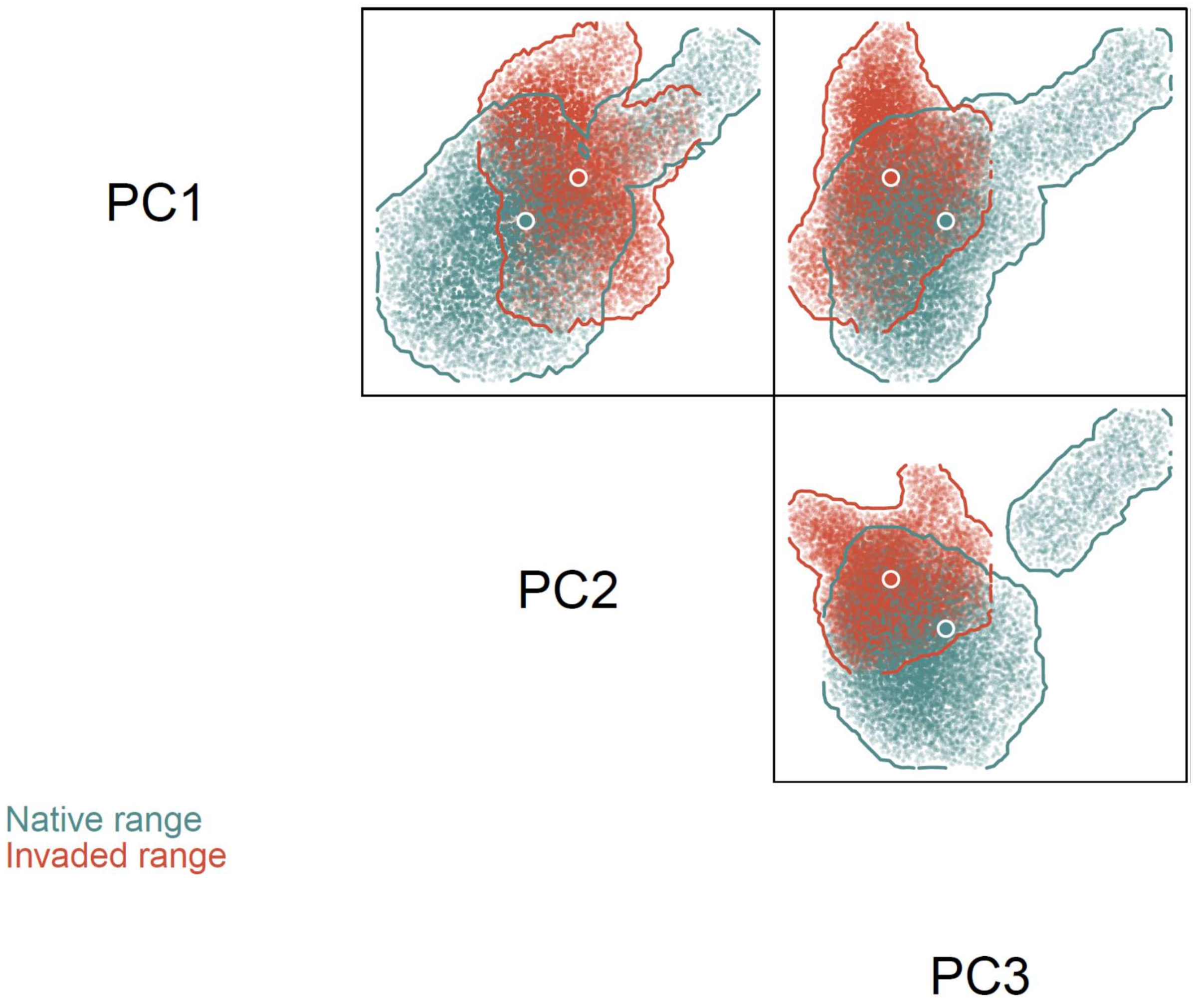


Fig. S6. Native and invasive niche hypervolumes of *Dikerogammarus haemobaphes*. The large dots represent niche centroids. The small dots are 10.000 random points sampled from each hypervolume to delineate its shape and boundary.

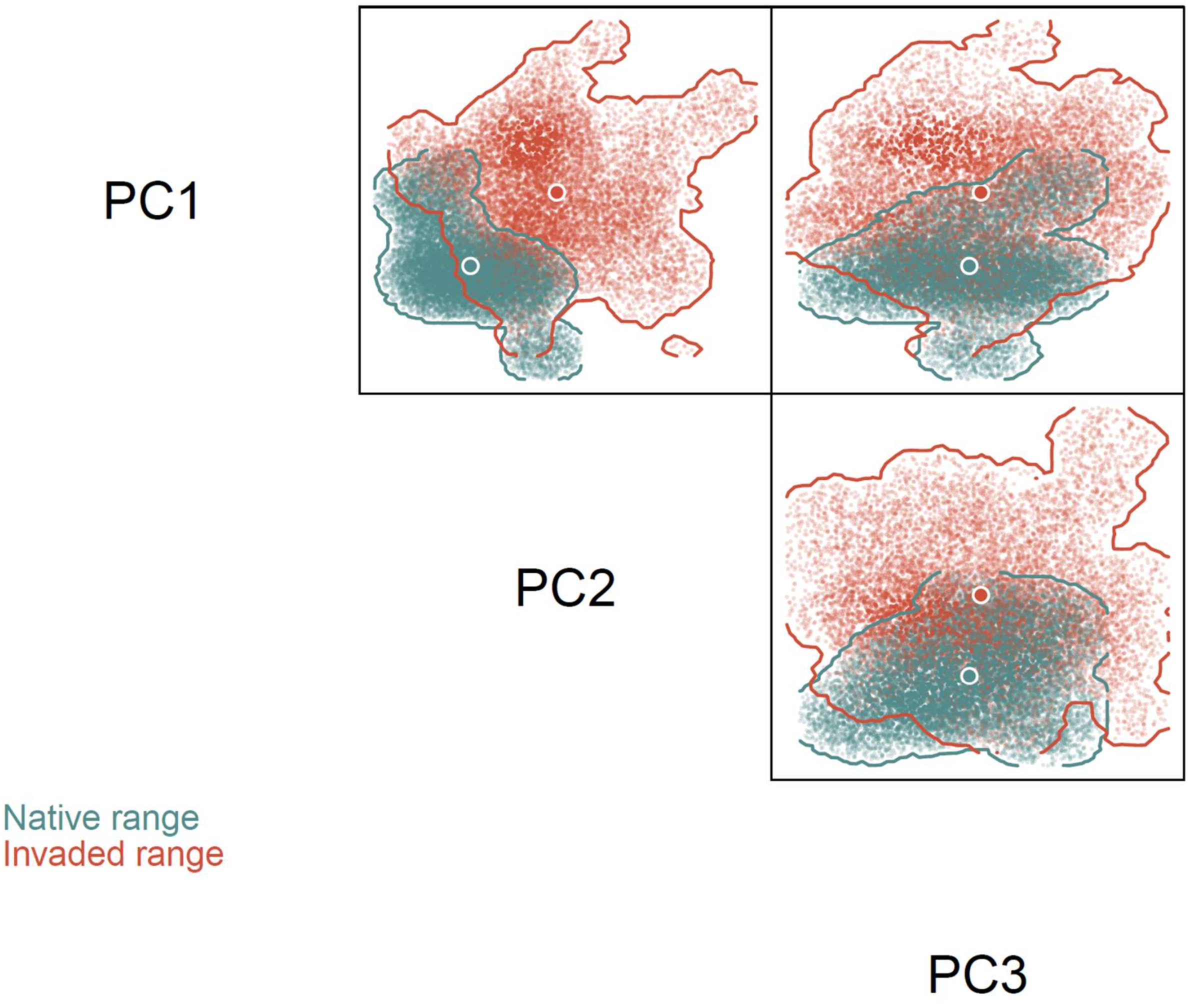


Fig. S7. Native and invasive niche hypervolumes of *Dikerogammarus villosus*. The large dots represent niche centroids. The small dots are 10.000 random points sampled from each hypervolume to delineate its shape and boundary.

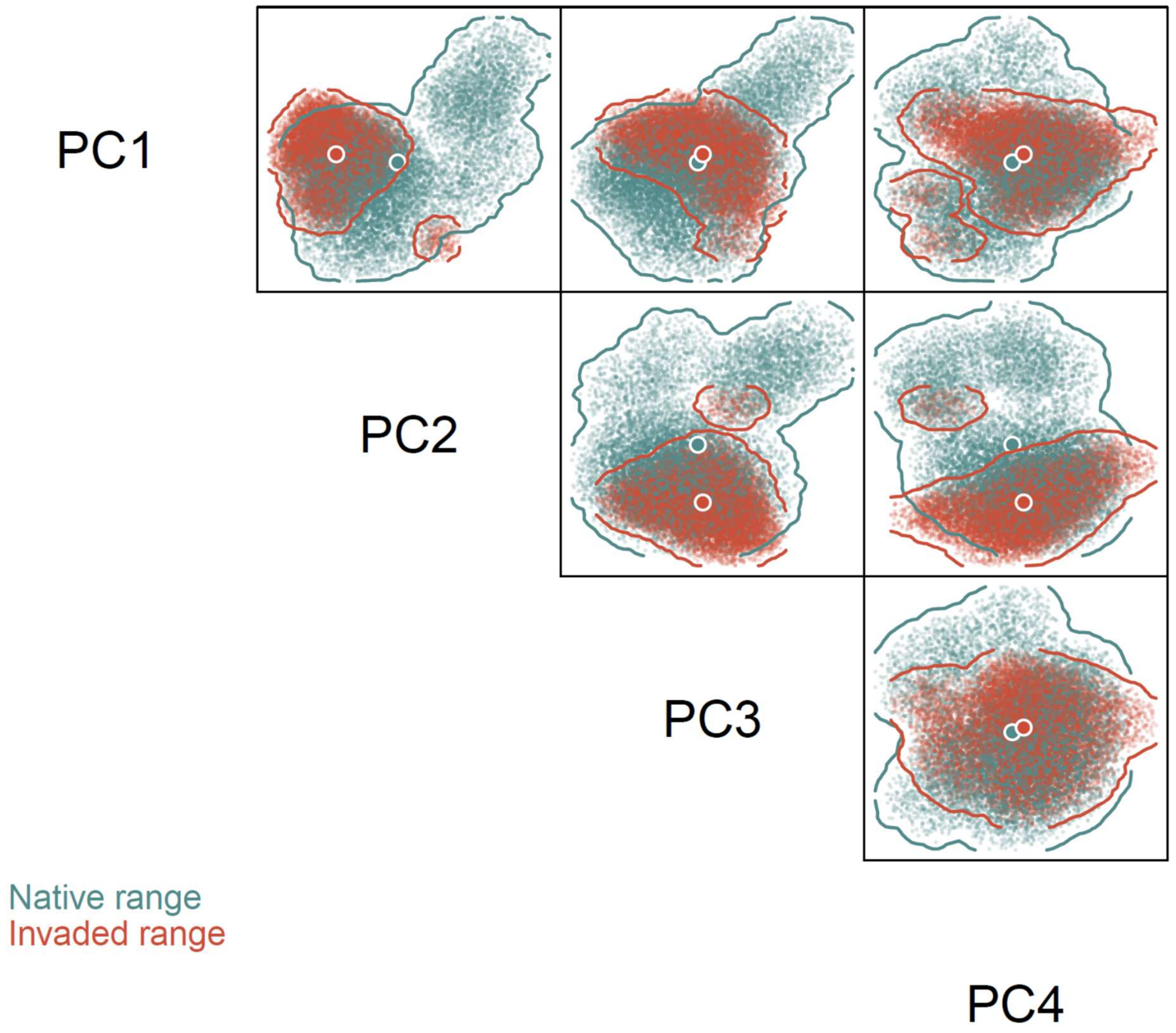


Fig. S8. Native and invasive niche hypervolumes of *Obesogammarus crassus*. The large dots represent niche centroids. The small dots are 10.000 random points sampled from each hypervolume to delineate its shape and boundary.

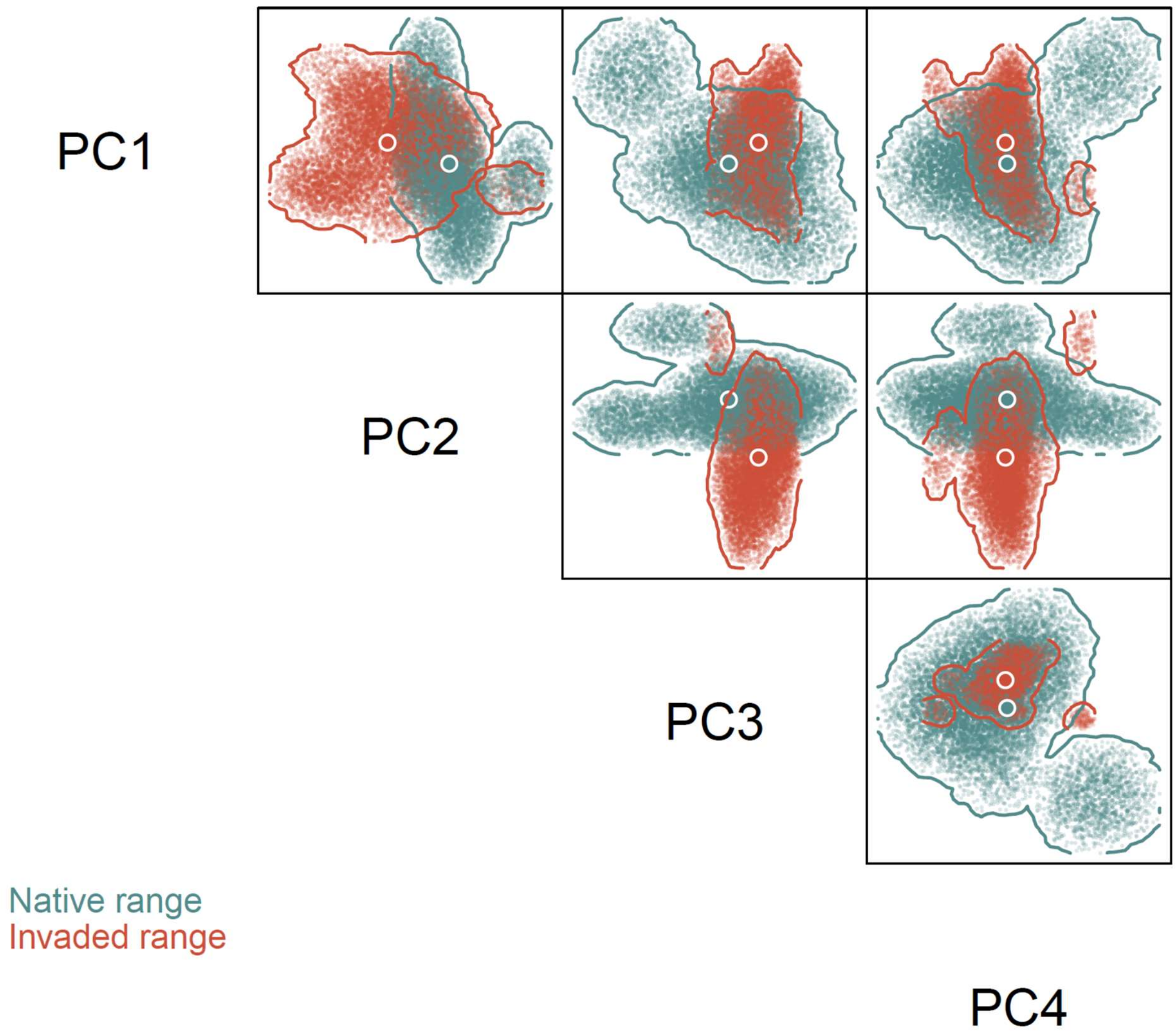


Fig. S9. Native and invasive niche hypervolumes of *Obesogammarus obesus*. The large dots represent niche centroids. The small dots are 10.000 random points sampled from each hypervolume to delineate its shape and boundary.

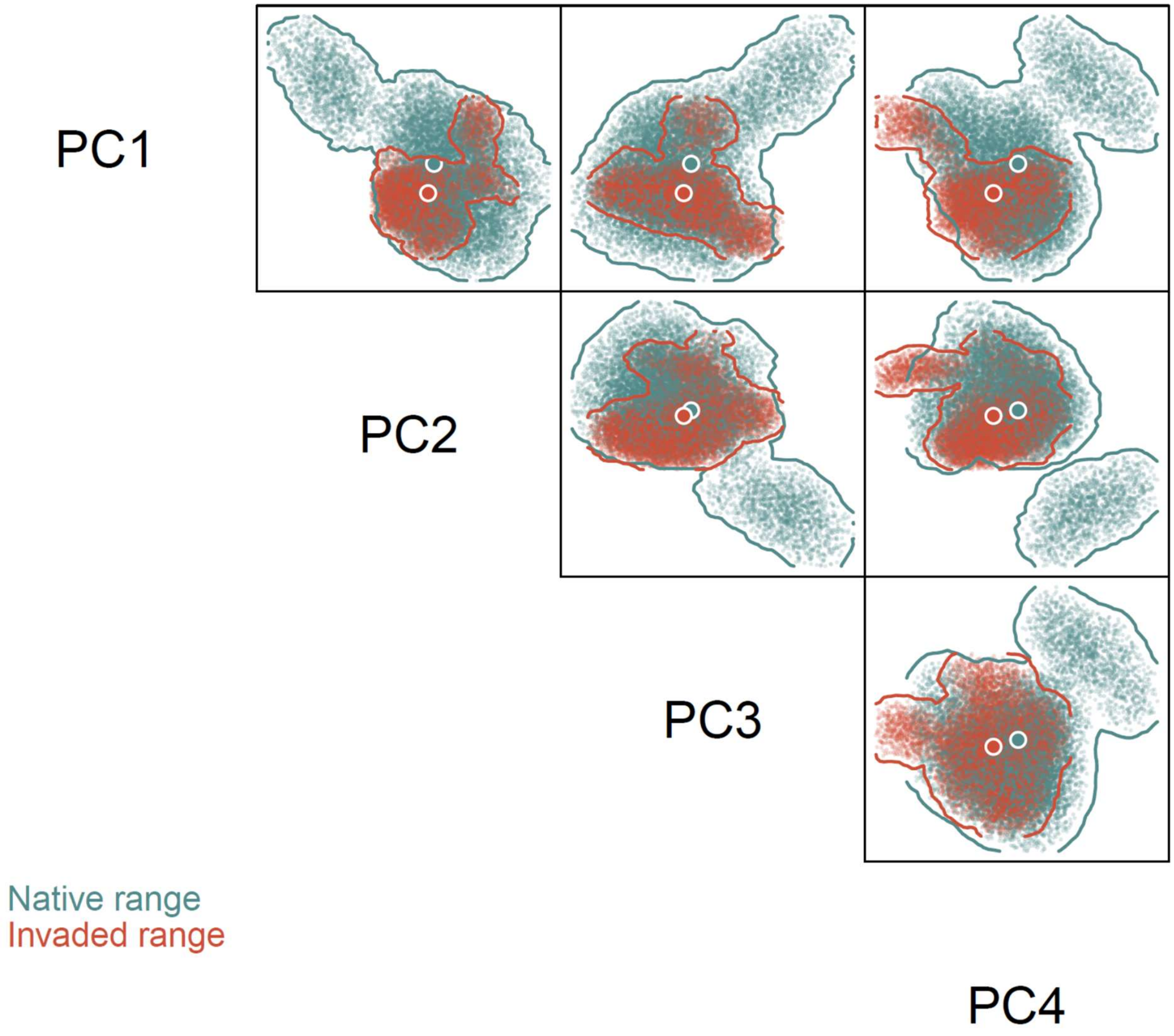


Fig. S10. Native and invasive niche hypervolumes of *Pontogammarus robustoides*. The large dots represent niche centroids. The small dots are 10.000 random points sampled from each hypervolume to delineate its shape and boundary.

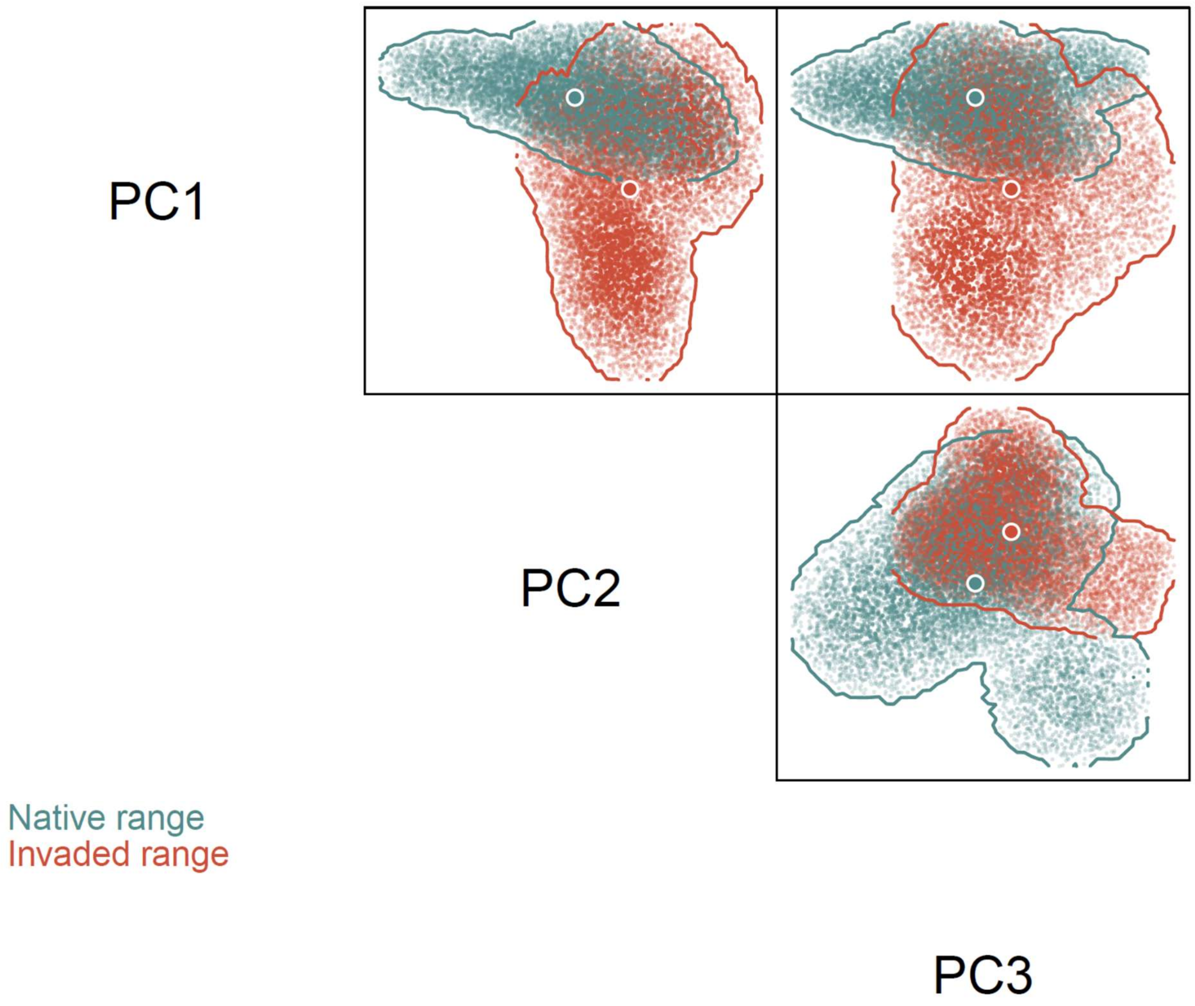


Fig. S11. Native and invasive niche hypervolumes of *Pontogammarus sarsi*. The large dots represent niche centroids. The small dots are 10.000 random points sampled from each hypervolume to delineate its shape and boundary.

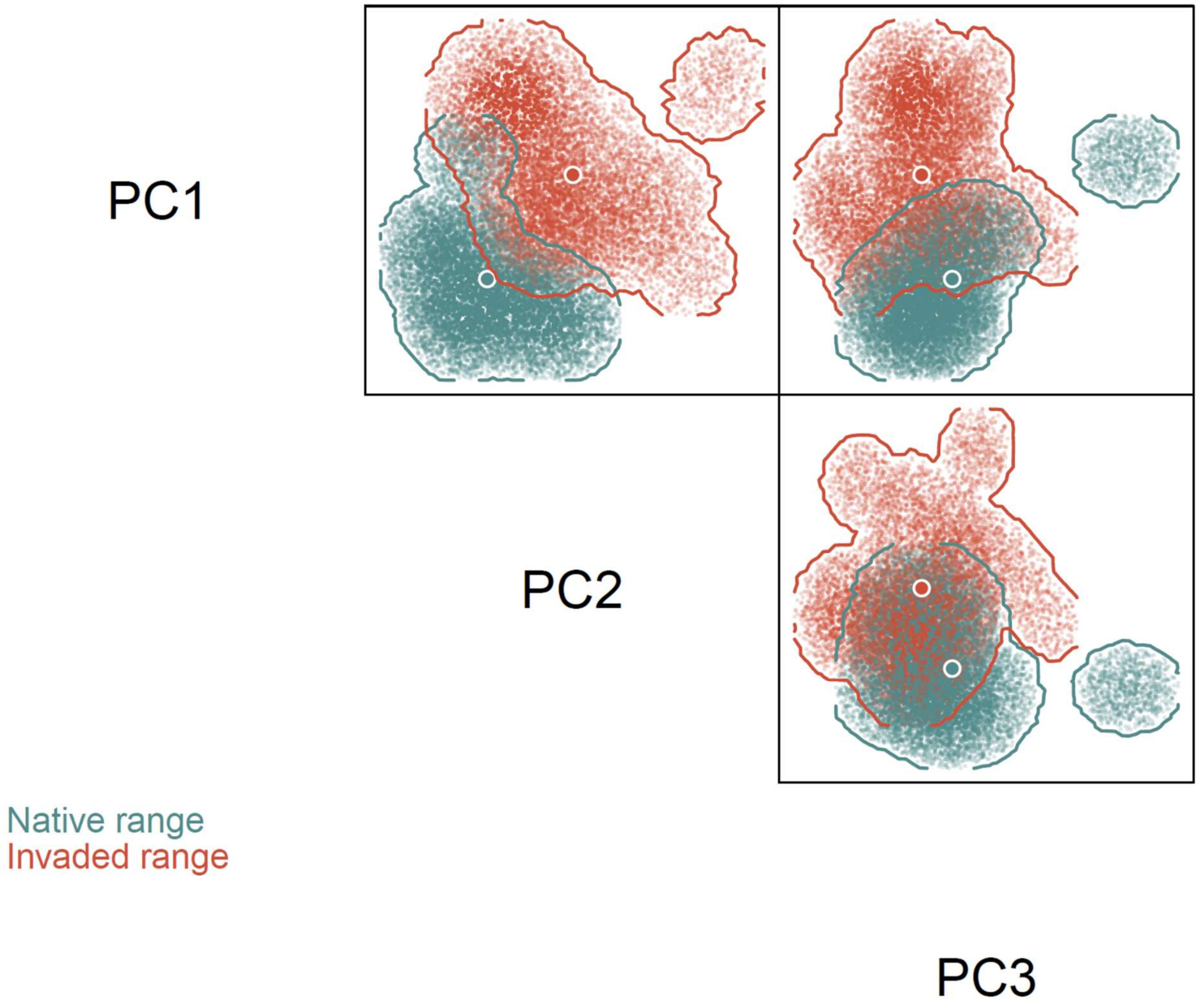


Fig. S12. Native and invasive niche hypervolumes of *Spirogammarus major*. The large dots represent niche centroids. The small dots are 10.000 random points sampled from each hypervolume to delineate its shape and boundary.
